# miRBind2 enables sequence-only prediction of miRNA binding and transcript repression

**DOI:** 10.64898/2026.03.19.712027

**Authors:** David Čechák, Dimosthenis Tzimotoudis, Stephanie Sammut, Katarina Gresova, Eva Marsalkova, David Farrugia, Panagiotis Alexiou

## Abstract

**Motivation:** MicroRNAs (miRNAs) regulate gene expression by guiding Argonaute proteins to partially complementary sites on target RNAs. While classical prediction methods rely on engineered features such as seed match categories, evolutionary conservation, and site context, recent advances in deep learning offer the potential to learn targeting rules directly from sequence. We developed a sequence-based deep learning model that improves miRNA target site prediction, and further validated the learned target site representations by extending the model to gene-level functional repression prediction.

**Results:** We introduce miRBind2, a deep learning method for miRNA target site prediction that incorporates a novel pairwise nucleotide representation capturing all possible miRNA-target nucleotide interactions, with a CNN-based architecture. miRBind2 outperforms previous SotA models across four independent datasets from the debiased miRBench benchmark, while using 92% fewer parameters. We show that the convolutional features and weights learned by miRBind2 can be transferred to transcript-level prediction by extending the miRBind2 architecture and fine-tuning it on miRNA perturbation experiments. This miRBind2-3UTR model predicts gene repression from sequence alone. On a dataset of 50,549 miRNA-gene pairs, miRBind2-3UTR significantly outperforms TargetScan. These results show that deep models pretrained on target site data can capture regulatory signals and predict functional repression without requiring conventional engineered biological features.

**Availability:** Models and source code are freely available via GitHub (https://github.com/BioGeMT/miRBind_2.0). A publicly available web-tool for novel predictions and visualization is available at : (https://huggingface.co/spaces/dimostzim/BioGeMT-miRBind2)

## 1 Introduction

MicroRNAs (miRNAs) are small regulatory RNAs that have been recognized for their diverse functions, notably as master regulators of gene expression. Since their initial characterization (Lee et al., 1993), miRNAs have been established as crucial guide sequences that can direct proteins of the Argonaute family (AGO) to bind to sites on target RNAs. They are essential during embryogenesis, tissue development, and for maintaining homeostasis in adults (Bernstein et al., 2003; Ivey and Srivastava, 2010). Dysregulation of miRNAs has been implicated in a range of diseases, including cancer (Di Martino et al., 2025; Jurj et al., 2026; Ortolano et al., 2026), cardiovascular diseases (van Rooij et al., 2006; Ikeda et al., 2007; Thum and Condorelli, 2015; Joglekar et al., 2025), neurological disorders (Hébert and De Strooper, 2009; Chrysanthou et al., 2025), and various immune conditions (O’Connell et al., 2007; Sonkoly and Pivarcsi, 2009; Dai and Ahmed, 2011; Prasad et al., 2025). They show great promise in clinical applications, holding potential as biomarkers (Condrat et al., 2020; Joglekar et al., 2025) and therapeutic agents (van Rooij and Olson, 2012; Rupaimoole and Slack, 2017; Yan et al., 2025), with recent inclusion in cutting-edge treatments like CAR-T therapy (Rad et al., 2022; Golinelli et al., 2025; Lonez et al., 2025), and synthetic biology (Abe et al., 2025).

The binding of AGO loaded with miRNAs to their target RNA is mediated by partial sequence complementarity between the “guide” miRNA and “target sites” most often found in the 3′ untranslated regions (3′UTRs) of the target RNA (Bartel, 2004). Such miRNA mediated binding between AGO and the target RNA leads to translational repression and can even promote the decay of the target RNA (Djuranovic et al., 2012; Jonas and Izaurralde, 2015; Bartel, 2018) (Figure 1A).

**Figure 1.**
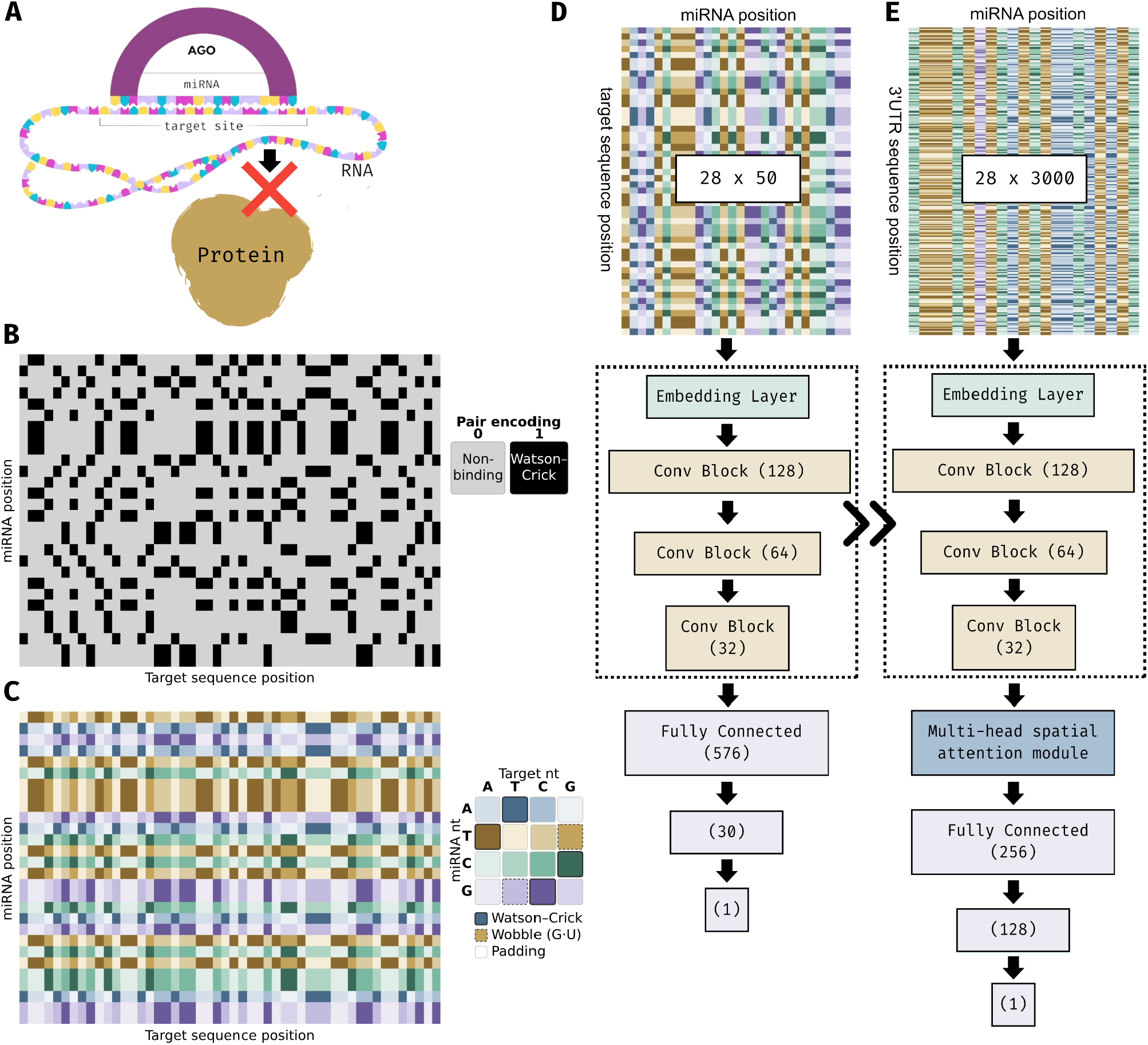
**(A)** miRNA mediated AGO binding to RNA targets leads to translational repression and degradation. **(B)** Binary representation of a miRNA-target pair. **(C)** Pairwise representation of a miRNA-target pair. **(D)** Simplified representation of the miRBind2 CNN model. **(E)** Simplified representation of the miRBind2-3UTR model

The rules of binding between a guide miRNA and its target sites are not fully understood. Most experimentally validated interactions between miRNAs and their targets involve complementarity concentrated in the “seed” region at the 5′ end of the miRNA sequence. The “canonical seed”, defined as a stretch of at least six fully Watson-Crick complementary nucleotides beginning at the second position from the 5′ end of the miRNA, has been the focus of most target site prediction tools to date (Lewis et al., 2005; Agarwal et al., 2015; Sammut et al., 2025). The stability of this interaction can be further enhanced by binding outside the seed area, commonly referred to as 3′ compensatory binding (Grimson et al., 2007; McGeary et al., 2022). However, target sites that do not rely on a “canonical seed” are also commonly found, with estimates of approximately 50% of target sites identified via high throughput sequencing lacking a canonical seed (Helwak et al., 2013; Klimentová et al., 2022; Hejret et al., 2023; Sammut et al., 2025).

Experimental identification of miRNA target sites requires significant resources and labor, and cannot easily scale to the full space of miRNA–target interactions, which leads to the need for in silico methods for miRNA target site prediction. This task can be defined as follows: given a miRNA sequence and a putative target site sequence, predict whether this miRNA would bind the presented target site.

For the task of miRNA target site prediction, we previously developed a sequence based deep learning model, miRBind (Klimentová et al., 2022), which at the time outperformed the state-of-the-art in this task. However, since then, we have reported evidence of a miRNA frequency class bias in commonly used miRNA:target-site datasets that inflated performance and hindered generalization in many existing algorithms including miRBind. We addressed this bias by constructing novel datasets, standardised miRNA target site prediction benchmarks, and retraining baseline models (Sammut et al., 2025). Specifically, a retrained Convolutional Neural Network (CNN) model with the architecture first proposed in miRBind (Klimentová et al., 2022) outperformed the state of the art in the corrected target site prediction benchmarks, and set the new baseline for the task of miRNA target site prediction (Sammut et al., 2025).

The task downstream of miRNA target site prediction is the prediction of the effect that a miRNA has on a target RNA as a whole. This ‘microRNA target gene prediction’ operates at the transcript level and is a related but separate task. Functional miRNA target prediction on 3′UTRs has a long history (Alexiou et al., 2009; Peterson et al., 2014; Hwang et al., 2023). State-of-the-art (SotA) approaches such as TargetScan (Agarwal et al., 2015) combine multiple signals beyond raw sequence, such as evolutionary conservation, site context, and additional engineered features to predict the effect of miRNAs on their target transcripts. These methods often employ simple representations of target sites such as seed length counting or categories, and then other features do the heavy lifting of prediction. The fundamental question is: could an improvement of target site prediction from sequence only, alleviate the need for other secondary engineered features and simplify functional target prediction?

In this study, we (i) introduce miRBind2, a methodology that can be used to model and predict miRNA target sites using a novel sequence representation, (ii) deploy the miRBind2 model in a user friendly web-tool, and (iii) show the benefits of transfer learning from miRNA target site datasets to a miRNA functional prediction dataset, leveraging the abundant miRNA target sites data to strengthen performance in the miRNA functional prediction.

## 2 Methods

### 2.1 MicroRNA Target Site Prediction

We use the miRBench Python package version 1.0.2 (Sammut et al., 2025) to access available datasets, encoders, and models for miRNA target site prediction. We have used the miRBenchCNN_Manakov (Sammut et al., 2025) and TargetScanCnn_McGeary2019 (McGeary et al., 2019) models without modifications.

#### 2.1.1 Data

We downloaded pre-established train-test splits from the miRBench datasets (v5) (Sammut et al., 2025) on Zenodo (https://zenodo.org/records/14501607). We used the Manakov2022 train set for training, and all available test sets for evaluation (Manakov2022 test & left-out, Hejret2023 test, Klimentova2022 test).

Each dataset contains labeled pairs of miRNA and target site in a 1:1 positive-to-negative ratio. The negative pairs were generated in the original study by pairing miRNAs in the positive class to alternate target sites from the same class, following the assumption that non-co-occurring miRNA-target site pairs represent non-binding interactions (Sammut et al., 2025).

#### 2.1.2 miRBench CNN

miRBenchCNN_Manakov (Sammut et al., 2025) is a CNN comprising six convolutional blocks and two dense blocks, using the architecture and data representation originally proposed in miRBind (Klimentová et al., 2022). The input is encoded as a 2D binary matrix representing Watson-Crick complementarity between 20 nucleotides from the miRNA 5′ end and 50 nucleotides of the target site, where complementary base pairs (AU, UA, GC, CG) are assigned 1 and all others 0 (Figure 1B). Shorter sequences are padded with N, and longer sequences are trimmed. Convolutional layers use 5 × 5 kernels with leaky ReLU activation, batch normalization, and 30% dropout; the final dense layer ends with a sigmoid activation, outputting the miRNA:target site binding probability. The model was trained on the Manakov2022 train set using the Adam optimizer (learning rate 0.00152).

#### 2.1.3 TargetScan CNN

TargetScanCnn_McGeary2019 (McGeary et al., 2019) is a CNN comprising two convolutional layers and two dense layers. A feature representation is produced by computing the outer product between one-hot encodings of the first 10 nucleotides of the miRNA and a 12-nucleotide segment from the candidate target site. During inference, the model evaluates every possible 12-nucleotide window within the target sequence and reports the maximum score among these windows as the final prediction. The model was trained on data derived from RNA Bind-n-Seq (RBNS) and mRNA-transfection experiments.

#### 2.1.4 miRBind2

We developed a pairwise encoding scheme to represent miRNA-target site interactions as discrete nucleotide pair combinations rather than binary complementarity scores. With four possible nucleotides for the miRNA and target (A, T, C, G), this yields 16 combinations. With the addition of a padding token, 17 unique combinations are defined. Each miRNA-target interaction can thus be encoded as a three-dimensional tensor of shape (miRNA length) × (target length) × (number of nucleotide pair combinations (17)), where each nucleotide pair is represented as a one-hot encoded vector of length 17. This tensor is passed through a learnable embedding layer that maps each 17-dimensional one-hot vector to a continuous vector representation of dimension d = 8. This embedding layer learns distributed representations that capture the binding affinity characteristics of each nucleotide pair, allowing the model to discover optimal representations for all pairs, including Watson-Crick pairs (e.g., A–U, G–C), wobble pairs (G–U), and mismatches. (Figure 1C). The input size of miRNA is 28 nt (the longest miRNA in the train set) and the input size of a target site is 50 nt. Longer sequences are cut, and shorter sequences are padded with N.

We trained a CNN model, using Bayesian optimization to optimize model hyperparameters and architecture. The hyperparameter search space included batch size (16–128), embedding dimension (2–16), learning rate (10^−5^–10^−2^, log scale), and dropout rate (0.1–0.5). The architecture search space included the number of convolutional layers (1–4), number of feature maps per layer (starting from 128–256 for layer 1, halving for subsequent layers, floored at 32) and kernel sizes (3–15, constrained to prevent feature map collapse). During optimization, we calculated maximum viable kernel sizes at each layer to ensure sufficient spatial dimensions were preserved for subsequent operations.

We maximized the validation area under precision recall curve (PR-AUC) using a median pruning strategy to terminate unpromising trials early. Each trial was trained with early stopping (patience=5) on a 90/10 train-validation split. The optimization process evaluated architectures that maintained adequate feature map dimensions through the network, automatically pruning configurations that would result in degenerate representations.

### 2.2 MicroRNA Target Gene Prediction

MicroRNA target gene prediction is framed as a regression task for training purposes: given a miRNA sequence and a gene’s 3’ UTR sequence, predict the log_2_ fold change in mRNA expression caused by the miRNA. Each miRNA-gene pair has a single continuous label: the measured log_2_FC from the microarray. More negative predictions indicate stronger predicted repression. For evaluation, we use both regression metrics (Pearson, Spearman, R^2^) and classification metrics (ROC-AUC, AP), binarizing the continuous regression labels at a threshold of log_2_FC < −0.05 to define repressed genes (Figure 2A) (see Section 2.3). This formulation evaluates the methods on their ability to both quantify the magnitude of miRNA-mediated regulation and classify genes as targets or non-targets.

**Figure 2:**
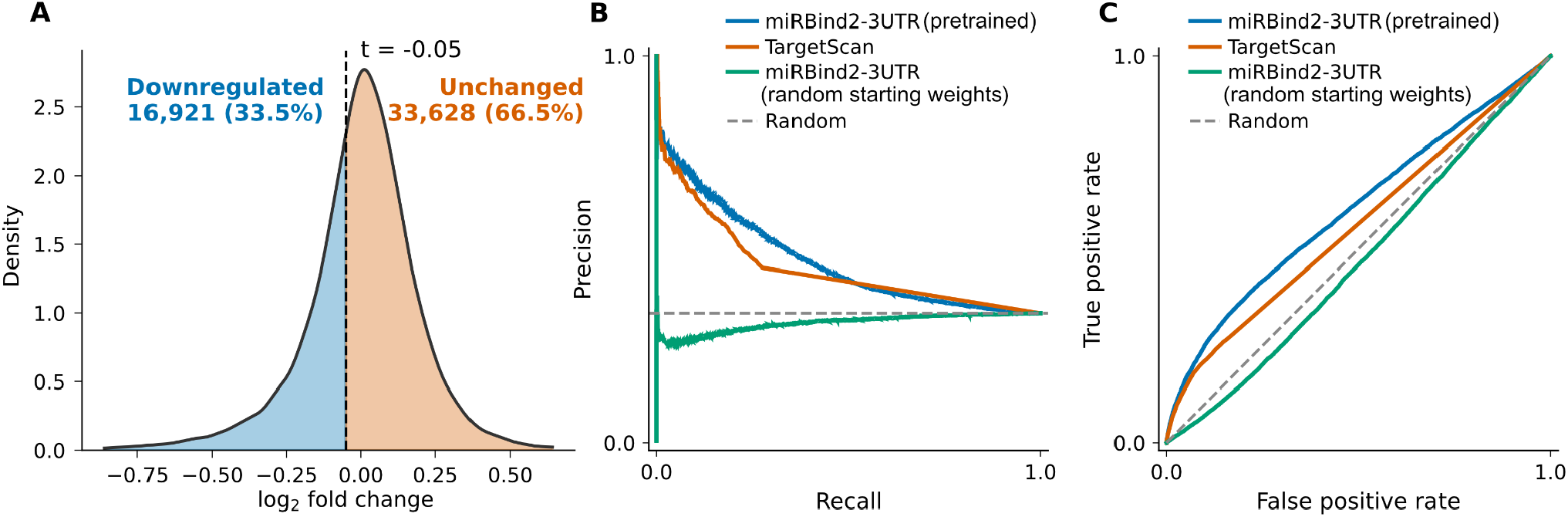
**(A)** Density distribution of measured log_2_ fold changes across miRNA–gene pairs in the test set. The dashed vertical line marks the classification threshold (log_2_FC < −0.05) used to binarize regression labels into downregulated (blue, 16,921 pairs, 33.5%) and unchanged (orange, 33,628 pairs, 66.5%) classes. (B) PR and (C) ROC curve on the functional targeting test set. The miRBind2-3UTR improvement over TargetScan (Agarwal et al., 2015) was statistically highly significant. A Williams’ test for dependent correlations (Steiger, 1980) yielded t = 204.3 (p ≈ 0) for the Pearson difference, and bootstrap 95% confidence intervals for the correlation difference excluded zero (ΔPearson = +0.062, 95% CI [+0.050, +0.073]; ΔSpearman = +0.043, 95% CI [+0.033, +0.052]). The ROC-AUC difference (0.603 vs. 0.561, binarized at log_2_FC < −0.05) was significant by DeLong’s test (DeLong et al., 1988; z = 16.3, p = 1.6 × 10^−59^).

#### 2.2.1 Data

For the train set, normalized mRNA fold-change values from a compendium of 74 small RNA transfection experiments in HeLa cells were taken from Supplementary File 1 of (Agarwal et al., 2015), retaining only the subset of genes indicated in Supplementary File 1 as passing the original authors’ filtering criteria for model training (i.e., genes with detectable expression and a dominant 3′UTR isoform in HeLa cells). Small RNA sequences for each experiment were obtained from the supplementary data of (Garcia et al., 2011). Training data were assembled following the methodology and supplementary materials of (Agarwal et al., 2015). Human 3′ UTR sequences were obtained from the published gene sequences (file: “human.3utrs.90%” (Agarwal et al., 2015)), which contain one representative 3′ UTR per gene, restricted to genes for which ≥90% of 3P-seq tags in HeLa cells corresponded to a single dominant isoform (Agarwal et al., 2015). 3’UTR sequences shorter than 25 nt were excluded. These 3P-seq-filtered UTR sequences are based on the hg19 (GRCh37) genome assembly. Each (3′ UTR, sRNA, fold-change) combination was assembled into a single training example. Extreme fold-change outliers beyond the 0.01st and 99.99th percentiles were removed to limit the influence of technical artifacts or aberrant measurements on model training, yielding a final dataset of 265,180 training examples.

We compiled a test set of 50,549 miRNA-gene pairs (7 miRNAs × 7,486 genes) by linking experimentally measured log_2_ fold changes from miRNA transfection microarray experiments (Agarwal et al., 2015), and human 3’ UTR sequences from the UCSC Genome Browser (hg19). Only transcripts present in TargetScan’s prediction files for at least one miRNA were kept, to ensure both methods were evaluated on the same gene set (see Section 2.3 for details on how TargetScan scores were obtained). For classification evaluation, continuous log_2_FC labels were binarized at a threshold of log_2_FC < −0.05 to define repressed genes (see Section 2.3).

Sequences in both datasets were capped at a maximum gene length of 3000 nt and a miRNA length of 28 nt, shorter sequences are padded, longer sequences are cut at the end.

#### 2.2.2 Model Architecture

To predict miRNA-mediated RNA repression at the gene level, we developed a transfer learning approach that repurposes the target site-level miRBind2-seq model for log2fold-change prediction from 3′ UTR sequences.

The gene-level architecture (named miRBind2-3UTR) consists of three primary components: a feature encoder, a multi-head spatial attention module, and a regression head. The feature encoder reuses the convolutional layers of the trained miRBind2, transferred into the gene-level architecture with their pretrained weights. This encoder, trained on miRNA-target site classification, provides learned representations of local binding features. The input to the gene-level model is encoded using the same pairwise nucleotide scheme as miRBind2 (Methods 2.1.4), producing a tensor of shape (miRNA length × 3′ UTR length × 17), where the 3′ UTR length replaces the fixed 50 nt target site window. The feature encoder architecture is identical to that of miRBind2; only the input dimensions change to accommodate full-length 3′ UTRs. Because full-length 3′UTRs produce larger feature maps than the fixed-size 50-nucleotide target site windows used during pretraining, we introduced a multi-head spatial attention module to aggregate convolutional features across the 2D spatial dimensions. This attention mechanism follows the attention-based pooling proposed by (Ilse et al. 2018), extended to a multi-head formulation by (Keshvarikhojasteh et al. 2024). Each attention head applies two linear projections with a Tanh nonlinearity in between to compute a scalar attention score at each spatial position of the feature map. These scores are normalized across all spatial positions using softmax, producing attention weights that indicate the relative importance of each position. The resulting attention weights are used to compute a weighted sum of the feature vectors across all spatial positions, aggregating the variable-size 2D feature map into a single fixed-length representation. This formulation allows the model to learn and focus on the most informative regions of the 3′ UTR. Eight independent attention heads each learn a different weighting pattern, and their outputs are combined via learnable head weights into a final pooled feature vector. Layer normalization is applied before and after the attention computation. Finally, the regression head processes the resulting 32-dimensional feature vector through two fully connected layers (256 → 128 → 1) with layer normalization, LeakyReLU activation, and 30% dropout.

#### 2.2.3 Transfer Learning and Training Strategy

A central feature of our approach is the use of transfer learning from the miRBind2 target site models to the gene-level repression task. We employed several strategies to effectively adapt the pretrained features. We used discriminative learning rates (Howard and Ruder, 2018) to preserve the fundamental binding patterns learned during site-level training. We applied a lower learning rate (LR) to the pretrained layers while using a higher base LR for the newly initialized attention and regression layers.

To emphasize the prediction of significant repression events in the dataset with low density of strongly downregulated 3’ UTRs, we use a Weighted Mean Squared Error (WMSE) loss function. This function assigns a higher weight (w=3.0) to samples where the actual fold change is below a threshold of -0.01, ensuring the model prioritizes learning strong repression over neutral interactions.

The gene-level model was trained using the Adam optimizer (base LR of 0.001, pretrained LR factor of 0.01). We used a batch size of 128 and trained for up to 300 epochs, employing early stopping basyed on the validation set Pearson correlation coefficient with a patience of 25 epochs.

### 2.3 Evaluation

For the task of miRNA target site prediction, we use the miRBench Python Package version 1.0.2 to encode the test sets and run inference for miRBenchCNN_Manakov and TargetScanCnn_McGeary2019 models to obtain predictions. We have made these predictions publicly available at https://zenodo.org/records/18682335.

For the task of miRNA target gene prediction, we obtained prediction scores and gene annotations from the TargetScan 8.0 data repository (www.targetscan.org/cgi-bin/targetscan/data_download.vert80.cgi). We merged two prediction files, Conserved site context++ scores and Nonconserved site context++ scores, to obtain scores for all predicted target sites. We used weighted context++ scores exclusively, as they consistently outperformed the unweighted context++ score across all metrics on the test set. Following the original publication, site-level weighted context++ scores were summed across all predicted binding sites per miRNA-gene pair. The fold-change data, which defines the gene set under evaluation, is annotated with both RefSeq transcript IDs and gene symbols. Because TargetScan indexes its predictions by Ensembl transcript ID rather than RefSeq ID, we used the gene symbol to bridge the two datasets, looking up each gene’s representative Ensembl transcript in TargetScan’s Gene_info file. Gene_info designates a single representative transcript per gene, defined as the transcript with the most 3P-seq–supported polyadenylation sites, reflecting the most commonly used 3′ UTR isoform (Agarwal et al., 2015). The representative Ensembl transcript ID was then used to retrieve the aggregated TargetScan score. This resolved 92.1% of genes. For the remaining 643 genes (7.9%) whose gene symbols were not found in Gene_info, scores were aggregated directly by gene symbol. TargetScan does not assign a score to miRNA-gene pairs lacking a predicted canonical target site, which accounts for 79.5% of miRNA-gene pairs in the test set. miRNA–gene pairs without any predicted weighted context++ score received a TargetScan score of zero.

We evaluated miRNA target sites classification models using ROC-AUC and Average Precision (AP). We use AP rather than the interpolated PR-AUC metric, as AP handles tied scores better, an important consideration since TargetScan assigns zero to many miRNA-gene pairs.

For the miRNA target gene regression models, we used Pearson and Spearman correlation coefficients between predicted and measured log_2_FC, and the coefficient of determination (R^2^). To also evaluate it using classification metrics, we converted the regression outputs into binary labels using a threshold of log_2_FC < −0.05 and computed ROC-AUC and AP.

We applied: (i) Williams’ test for dependent correlations (Steiger, 1980), which accounts for the correlation between the two predictors when comparing their correlations with the same ground truth; (ii) bootstrap 95% confidence intervals on the Pearson and Spearman correlation differences (10,000 resamples; (Efron et al., 1994)); (iii) DeLong’s test for comparing ROC-AUC values from the same samples (DeLong et al., 1988); and (iv) bootstrap 95% confidence intervals on the AP difference, as no closed-form analogue of DeLong’s test exists for AP. Tests (iii) and (iv) were also used to evaluate the miRNA target site prediction models.

### 2.4 Explainability of model predictions

To identify which input positions drive the miRBind2 model’s predictions, we computed per-nucleotide attribution scores using GradientSHAP (Erion et al., 2021), a gradient-based approximation of Shapley values that satisfies key interpretability axioms including completeness and implementation invariance. GradientSHAP extends Integrated Gradients (Sundararajan et al., 2017) by averaging over a distribution of reference inputs rather than a single baseline, yielding more stable attribution estimates. We used the implementation provided by the Captum library (Kokhlikyan et al., 2020) with a zero baseline and 25 stochastic samples per input. Attributions were computed on the pairwise one-hot encoded input tensor of shape (miRNA length × target site length × 17), then summed along the pair-encoding axis to produce a two-dimensional interaction attribution map of shape (miRNA length × target site length) for each sample. We further reduced this two-dimensional map to one-dimensional profiles along each axis: one along the target site, obtained by taking the maximum absolute attribution across miRNA positions at each target position, and one along the miRNA, obtained analogously by taking the maximum absolute attribution across target positions at each miRNA position. Per-nucleotide interactions between miRNA and a target site were computed using an attribution sequence alignment algorithm from (Gresova et al. 2023) with the two-dimensional interaction attribution map of shape (miRNA length x target site length) as an input. To adapt the algorithm to pairwise input encoding, we have removed following steps from the original attribution sequence alignment algorithm: 1. remove negative scores from the score matrix, and 2. swap sign of scores for the mismatch positions in the scoring matrix.

## 3 Results and Discussion

### 3.1 miRBind2 improves performance on miRNA target site prediction task

In our previous work (Sammut et al., 2025), we introduced a CNN for miRNA target-site classification using a binary interaction representation of 20 nucleotides from the miRNA 5’ end and 50 nucleotides of the predicted target site. That model was meant to be an intentionally simple baseline, and was not subjected to systematic hyperparameter tuning. Here, we revisit this task with the goal of establishing a stronger and better-controlled model trained on the Manakov2022 debiased benchmark (Sammut et al., 2025).

First, we extend the binary encoding to a pairwise nucleotide representation which encodes more information beyond the canonical Watson-Crick binding patterns, that might be biologically relevant for the miRNA-target site interaction. We additionally extend the 20-nt miRNA length in the binary encoding to use the whole miRNA length, padding shorter sequences. This ensures there is no data loss.

Following the hyperparameter optimization described in the Methods (Section 2.1.4), we found that the best-performing architecture consists of an embedding layer (dimension 8), followed by three convolutional blocks with decreasing feature map counts (128, 64, 32). The three convolutional layers used kernel sizes 6×6, 3×3, and 3×3 with padding of 2, 1, and 1, respectively. Each convolutional block included batch normalization, 2×2 max pooling, and 20% dropout. The convolutional features were flattened to 576 dimensions and passed through two fully connected layers (576 → 30 → 1) with batch normalization and dropout after the first dense layer.

The optimised model with pairwise encoding had 92% less trainable parameters (147,241 compared to 1,863,617 in miRBenchCNN_Manakov), while maintaining the ability to solve the same task.

Benchmarking against the miRBenchCNN_Manakov model (Sammut et al., 2025) and the TargetScanCnn_McGeary2019 model (McGeary et al., 2019) revealed moderate but consistent performance improvement (Table 1, Table 2). Since miRBenchCNN substantially outperformed TargetScanCnn across all four datasets (e.g. ROC-AUC 0.81 vs 0.72 on Manakov Test), we focus on the comparison between miRBind2 and miRBenchCNN. miRBind2 achieved higher AP and ROC-AUC than miRBenchCNN on all four benchmarks (Tables 1 and 2). On the Manakov leftout set (20,054 pairs, balanced classes, miRNAs with no samples present in the training set), miRBind2 achieved ROC-AUC 0.81 versus 0.79 for miRBenchCNN, a highly significant difference (DeLong z = 11.13, p = 9.1 × 10^−29^; bootstrap 95% CI on the difference: [+0.018, +0.025]). AP on the same set was 0.83 versus 0.81 (95% CI [+0.018, +0.025]). On the larger Manakov test set (327,129 pairs), miRBind2 achieved ROC-AUC 0.82 versus 0.81 for miRBenchCNN (DeLong z = 33.84, p = 4.5 × 10^−251^; 95% CI [+0.011, +0.013]) and AP 0.85 versus 0.84 for miRBenchCNN (95% CI [+0.009, +0.010]); both differences were significant, with confidence intervals that exclude zero. The gains were largest on the Hejret dataset (AP 0.86 vs 0.84, ROC-AUC 0.84 vs 0.83) and the Klimentova dataset (AP 0.86 vs 0.84, ROC-AUC 0.83 vs 0.80), which use independent AGO-CLIP protocols, suggesting that miRBind2 generalises across experimental contexts rather than reflecting overfitting to a single data source.

**Table 1.** Precision-Recall, Average Precision Score for 4 benchmarking datasets.

|  | TargetScanCnn<br>McGeary2019 | miRBenchCNN_<br>Manakov | miRBind2 |
| --- | --- | --- | --- |
| Manakov Test | 0.77 | 0.84 | **0.85** |
| Manakov Leftout | 0.76 | 0.81 | **0.83** |
| Hejret Test | 0.74 | 0.84 | **0.86** |
| Klimentova Test | 0.71 | 0.84 | **0.86** |

**Table 2.** AU-ROC for 4 benchmarking datasets.

|  | TargetScanCnn<br>McGeary2019 | miRBenchCNN_<br>Manakov | miRBind2 |
| --- | --- | --- | --- |
| Manakov Test | 0.72 | 0.81 | **0.82** |
| Manakov Leftout | 0.72 | 0.79 | **0.81** |
| Hejret Test | 0.67 | 0.83 | **0.84** |
| Klimentova Test | 0.71 | 0.80 | **0.83** |

### 3.2 Target-site transfer learning improves 3′UTR-level functional prediction

To test whether target site learning transfers to downstream functional effects, we used our target site predictor miRBind2 as a pretrained backbone for 3′UTR-level functional prediction, fine-tuning it on miRNA perturbation experiments (Figure 1E). The task is framed as regression: given a miRNA sequence and a gene’s full 3′UTR, predict the log_2_ fold change in RNA expression following miRNA transfection. Each miRNA-gene pair carries a single continuous label derived from microarray measurements, where more negative values indicate stronger repression. The test set comprised 50,549 miRNA-gene pairs (7 miRNAs × 7,486 genes), of which 33.5% showed downregulation (log_2_FC < −0.05), enabling evaluation under both regression and classification frameworks (Figure 2A).

We extended the pairwise nucleotide representation to accommodate 3′UTRs of length up to 3,000 nt and transferred the trained convolutional layers from miRBind2 into a gene-level architecture (miRBind2-3UTR) augmented with multi-head spatial attention and a regression head (see Methods 2.2.2).

Pretraining on target site data provided a substantial benefit. The same architecture trained from random initialisation performed near or below chance across all metrics (AP 0.31, ROC-AUC 0.47, Pearson −0.02), whereas fine-tuning from binding-site pretraining yielded markedly stronger performance (Table 3), with precision-recall and ROC curves showing clear separation from the non-pretrained model (Figures 2B and 2C). Freezing the pretrained convolutional layers for the first 30 epochs of fine-tuning did not yield additional gains over end-to-end training from the start, suggesting that full adaptation of the pretrained features to the gene-level task is beneficial under our current setup.

**Table 3.** Regression and classification metrics on the functional targeting test set.

| Model | Regression Task |  |  | Classification Task |  |
| --- | --- | --- | --- | --- | --- |
|  | Pearson | Spearman | R<sup>2</sup> | APS | AU-ROC |
| miRBind2-3UTR (random starting weights) | -0.02 | -0.02 | -0.05 | 0.31 | 0.47 |
| miRBind2-3UTR (pretrained) | **0.30** | **0.20** | **0.07** | **0.47** | **0.60** |
| TargetScan | 0.24 | 0.15 | 0.04 | 0.41 | 0.56 |

We compared miRBind2-3UTR, which operates on raw sequence alone, against TargetScan’s weighted context++ score (Agarwal et al., 2015), a model that incorporates multiple engineered features beyond sequence, such as evolutionary conservation, target site accessibility, and seed-match types. Despite relying solely on sequence, miRBind2-3UTR outperformed TargetScan (Agarwal et al., 2015) across all metrics. Under regression, miRBind2-3UTR achieved a Pearson correlation of 0.30 with measured fold changes compared to 0.24 for TargetScan (Spearman 0.19 vs 0.15; R^2^ 0.07 vs 0.04). The Pearson difference was highly significant (Williams’ test t = 204.33, p ≈ 0; bootstrap 95% CI [+0.050, +0.073]), as was the Spearman difference (95% CI [+0.033, +0.052]). Under classification (log_2_FC < −0.05), miRBind2-3UTR achieved ROC-AUC 0.60 versus 0.56 for TargetScan (DeLong z = 16.27, p = 1.6 × 10^−59^; 95% CI [+0.037, +0.047]) and AP 0.47 versus 0.41 (95% CI [+0.055, +0.066]). The advantage was consistent across all seven miRNAs, with per-miRNA Pearson differences ranging from +0.01 (miR-215-5p, Williams’ p = 1.3 × 10^−109^) to +0.10 (miR-106b-5p, p ≈ 0). Per-miRNA ROC-AUC differences were significant for six of seven miRNAs (DeLong p < 10^−7^), except for miR-215-5p (p = 0.32).

These results are notable given that TargetScan draws on evolutionary conservation, target site accessibility, and seed-match types (Agarwal et al., 2015), annotations that are unavailable for synthetic miRNAs or UTRs, non-model organisms, or novel transcripts, where a sequence-only model such as miRBind2-3UTR avoids that limitation entirely. Moreover, TargetScan does not assign scores to miRNA-gene pairs lacking a predicted canonical target site, which accounts for the large majority of pairs in the test set, leaving most of the transcriptome undifferentiated. Because miRBind2-3UTR operates directly on sequence, it produces graded predictions for all pairs, which is particularly relevant given that an estimated 50% of target sites identified via high-throughput sequencing lack a canonical seed (Helwak et al., 2013; Klimentová et al., 2022; Hejret et al., 2023; Sammut et al., 2025). The convolutional features learned during the target-site prediction task carry over information relevant to functional repression, not only binding, supporting the value of transfer learning between these two related tasks. These findings do not diminish the utility of curated biophysical features; they show that transfer learning from binding-site data enhances 3′UTR-level functional prediction from sequence, establish sequence-derived representations as a strong standalone foundation, and open the door to future hybrid approaches that combine data-driven and knowledge-driven signals.

### 3.3 Web-based tool for miRNA target site prediction

To make miRBind2 accessible to the broader community, we developed an interactive web-based tool. The interface supports single target queries, in which a user submits one miRNA and one target-site sequence, as well as bulk mode, in which many pairs can be uploaded simultaneously. For each submitted pair, the tool returns the predicted binding probability and displays an interaction attribution map derived from GradientSHAP (see Methods 2.4), allowing the user to visually inspect which nucleotide positions contribute most to the prediction. If the user provides ground-truth labels alongside their input, the tool additionally computes classification metrics (ROC-AUC, AP) for the submitted batch, enabling rapid benchmarking on custom datasets without requiring local installation or computational resources.

## Acknowledgements

We would like to thank Prof. Jean-Paul Ebejer, and Dr. Vlastimil Martinek for discussions during the analysis of data and preparation of this manuscript.

## Data availability

Models and source code are freely available via GitHub (https://github.com/BioGeMT/miRBind_2.0).

A publicly available web-tool for novel predictions is available at : (https://huggingface.co/spaces/dimostzim/BioGeMT-miRBind2).

## References

Abe,I. et al. (2025) Split RNA switch orchestrates pre-and post-translational control to enable cell type-specific gene expression. Nature Communications, 16, 5362.

Agarwal,V. et al. (2015) Predicting effective microRNA target sites in mammalian mRNAs. Elife, 4.

Alexiou,P. et al. (2009) Lost in translation: an assessment and perspective for computational microRNA target identification. Bioinformatics, 25, 3049–3055.

Bartel,D.P. (2018) Metazoan MicroRNAs. Cell, 173, 20–51.

Bartel,D.P. (2004) MicroRNAs: genomics, biogenesis, mechanism, and function. Cell, 116, 281–297.

Bernstein,E. et al. (2003) Dicer is essential for mouse development. Nat. Genet., 35, 215–217.

Chrysanthou,M. et al. (2025) The Role of miRNAs in Parkinson’s Disease: A Systematic Review. Int J Mol Sci, 26.

Condrat,C.E. et al. (2020) miRNAs as Biomarkers in Disease: Latest Findings Regarding Their Role in Diagnosis and Prognosis. Cells, 9.

Dai,R. and Ahmed,S.A. (2011) MicroRNA, a new paradigm for understanding immunoregulation, inflammation, and autoimmune diseases. Transl. Res., 157, 163–179.

DeLong,E.R. et al. (1988) Comparing the areas under two or more correlated receiver operating characteristic curves: a nonparametric approach. Biometrics, 44, 837–845.

Di Martino,M.T. et al. (2025) MicroRNA in cancer therapy: breakthroughs and challenges in early clinical applications. J Exp Clin Cancer Res, 44, 126.

Djuranovic,S. et al. (2012) miRNA-mediated gene silencing by translational repression followed by mRNA deadenylation and decay. Science, 336, 237–240.

Efron,B. et al. (1994) An introduction to the bootstrap Chapman & Hall/CRC, Philadelphia, PA.

Erion,G. et al. (2021) Improving performance of deep learning models with axiomatic attribution priors and expected gradients. Nat. Mach. Intell., 3, 620–631.

Garcia,D.M. et al. (2011) Weak seed-pairing stability and high target-site abundance decrease the proficiency of lsy-6 and other microRNAs. Nat Struct Mol Biol, 18, 1139–1146.

Grešová,K. et al. (2023) Using Attribution Sequence Alignment to Interpret Deep Learning Models for miRNA Binding Site Prediction. Biology, 12.

Golinelli,G. et al. (2025) Multiplex engineering using microRNA-mediated gene silencing in CAR T cells. Front Immunol, 16, 1647433.

Grimson,A. et al. (2007) MicroRNA targeting specificity in mammals: determinants beyond seed pairing. Mol Cell, 27, 91–105.

Hébert,S.S. and De Strooper,B. (2009) Alterations of the microRNA network cause neurodegenerative disease. Trends Neurosci., 32, 199–206.

Hejret,V. et al. (2023) Analysis of chimeric reads characterises the diverse targetome of AGO2-mediated regulation. Sci. Rep., 13, 22895.

Helwak,A. et al. (2013) Mapping the human miRNA interactome by CLASH reveals frequent noncanonical binding. Cell, 153, 654–665.

Howard,J. and Ruder,S. (2018) Universal language model fine-tuning for text classification. In, Proceedings of the 56th Annual Meeting of the Association for Computational Linguistics (Volume 1: Long Papers). Association for Computational Linguistics, Stroudsburg, PA, USA.

Hwang,H. et al. (2023) Determinants of Functional MicroRNA Targeting. Mol Cells, 46, 21–32.

Ikeda,S. et al. (2007) Altered microRNA expression in human heart disease. Physiol. Genomics, 31, 367–373.

Ilse,M. et al. (2018) Attention-based deep multiple instance learning. arXiv [cs.LG].

Ivey,K.N. and Srivastava,D.(2010) MicroRNAs as regulators of differentiation and cell fate decisions. Cell Stem Cell, 7, 36–41.

Joglekar,M.V. et al. (2025) A microRNA-based dynamic risk score for type 1 diabetes. Nat Med, 31, 2622–2631.

Jonas,S. and Izaurralde,E. (2015) Towards a molecular understanding of microRNA-mediated gene silencing. Nat Rev Genet, 16, 421–433.

Jurj,A. et al. (2026) MicroRNAs in oncology: a translational perspective in the era of AI. Nature Reviews Clinical Oncology, 1–21.

Keshvarikhojasteh,H. et al. (2024) Multi-head attention-based Deep Multiple Instance Learning. arXiv [cs.CV].

Klimentová,E. et al. (2022) miRBind: A deep learning method for miRNA binding classification. Genes, 13, 2323.

Kokhlikyan,N. et al. (2020) Captum: A unified and generic model interpretability library for PyTorch. arXiv [cs.LG].

Lee,R.C. et al. (1993) The C. elegans heterochronic gene lin-4 encodes small RNAs with antisense complementarity to lin-14. Cell, 75, 843–854.

Lewis,B.P. et al. (2005) Conserved seed pairing, often flanked by adenosines, indicates that thousands of human genes are microRNA targets. Cell, 120, 15–20.

Lonez,C. et al. (2025) Clinical Proof-of-Concept of a Non-Gene Editing Technology Using miRNA-Based shRNA to Engineer Allogeneic CAR T-Cells. Int J Mol Sci, 26.

McGeary,S.E. et al. (2022) MicroRNA 3’-compensatory pairing occurs through two binding modes, with affinity shaped by nucleotide identity and position. Elife, 11.

McGeary,S.E. et al. (2019) The biochemical basis of microRNA targeting efficacy. Science, 366.

O’Connell,R.M. et al. (2007) MicroRNA-155 is induced during the macrophage inflammatory response. Proc. Natl. Acad. Sci. U. S. A., 104, 1604–1609.

Ortolano,A. et al. (2026) miRNA-driven cancer cell plasticity, tolerance and therapy resistance: lessons from melanoma. Mol Cancer.

Peterson,S.M. et al. (2014) Common features of microRNA target prediction tools. Front Genet, 5, 23.

Prasad,S. et al. (2025) The role of microRNAs and long non-coding RNAs in epigenetic regulation of T cells: implications for autoimmunity. Front Immunol, 16, 1695894.

Rad,S.M.A.H. et al. (2022) MicroRNA-mediated metabolic reprogramming of chimeric antigen receptor T cells. Immunol. Cell Biol., 100, 424–439.

van Rooij,E. et al. (2006) A signature pattern of stress-responsive microRNAs that can evoke cardiac hypertrophy and heart failure. Proc. Natl. Acad. Sci. U. S. A., 103, 18255–18260.

van Rooij,E. and Olson,E.N. (2012) MicroRNA therapeutics for cardiovascular disease: opportunities and obstacles. Nat. Rev. Drug Discov., 11, 860–872.

Rupaimoole,R. and Slack,F.J. (2017) MicroRNA therapeutics: towards a new era for the management of cancer and other diseases. Nat. Rev. Drug Discov., 16, 203–222.

Sammut,S. et al. (2025) miRBench: novel benchmark datasets for microRNA binding site prediction that mitigate against prevalent microRNA frequency class bias. Bioinformatics, 41, i542–i551.

Sonkoly,E. and Pivarcsi,A. (2009) Advances in microRNAs: implications for immunity and inflammatory diseases. J. Cell. Mol. Med., 13, 24–38.

Steiger,J.H. (1980) Tests for comparing elements of a correlation matrix. Psychol. Bull., 87, 245–251.

Sundararajan,M. et al. (2017) Axiomatic attribution for deep networks. arXiv [cs.LG].

Thum,T. and Condorelli,G. (2015) Long noncoding RNAs and microRNAs in cardiovascular pathophysiology. Circ. Res., 116, 751–762.

Yan,Y. et al. (2025) Advances in RNA-based cancer therapeutics: pre-clinical and clinical implications. Mol Cancer, 24, 251.

